# Elimination of ribosome inactivating factors improves the efficiency of *Bacillus subtilis and Saccharomyces cerevisiae* cell-free translational systems

**DOI:** 10.1101/424044

**Authors:** Tetiana Brodiazhenko, Marcus Johansson, Hiraku Takada, Tracy Nissan, Vasili Hauryliuk, Victoriia Murina

**Author notes:** Contact details of corresponding authors:Victoriia Murina, Vasili Hauryliuk.

## Abstract

Cell-free translational systems based on cellular lysates optimized for *in vitro* protein synthesis have multiple applications both in basic and applied science, ranging from studies of translation regulation to cell-free production of proteins and ribosome-nascent chain complexes. In order to achieve both high activity and reproducibility in a translational system, it is essential that the ribosomes in the cellular lysate are enzymatically active. Here we demonstrate that genomic disruption of genes encoding ribosome inactivating factors – HPF in *Bacillus subtilis* and Stm1 in *Saccharomyces cerevisiae* – robustly improve the activities of bacterial and yeast translational systems. A possible next step in developing strains for production of even more efficient cell-free translation systems could be achieved by combining a genomic disruption of the ribosome hibernation machinery with inactivation of the genes responsible for proteolysis and RNA degradation.

## Introduction

Cell-free translational systems based on cellular lysates optimized for *in vitro* protein synthesis have multiple applications both in basic and applied science, ranging from studies of translation regulation^1^ to cell-free production of recombinant proteins^2^ and ribosome-nascent chain complexes^3^. The preparation of cell-free translational systems is a compromise between, on one hand, the desired properties, such as high synthetic activity and reproducibility of the system and, on the other hand, simplicity of generating robust extracts as well as economic considerations. In laboratory settings, the most convenient and readily accessible method of producing biomass is by growing cells in a batch format in flasks. In this case, large-scale production of highly translationally active exponentially growing cells can be challenging due to relatively low yields. To maximize extract yields, one can harvest cultures in late exponential / early stationary phase. While this provides more biomass, there is a drawback in that cells often reduce their translational capacity during slow growth^4^. Importantly, the translational activity decreases upon exiting rapid exponential growth – and an important mechanism at play is the reduction of the active ribosomal concentration via ribosomal sequestration into inactive complexes by dedicated regulatory protein factors. Bacteria reduce their translational capacity by forming inactive ribosome dimers, so-called 100S ribosomes^5,6^. In Gamma-proteobacteria, such as *Escherichia coli*, this process is mediated by two cooperating factors: the Hibernation Promoting Factor (HPF) and the Ribosome Modulation Factor (RMF)^7^. In the majority of bacterial species, 100S formation is mediated by one factor – HPF – the long version of the ‘short’ HPF present in the Gamma-proteobacteria^8-10^. In budding yeast, the Stm1 protein acts as a translational repressor^11^ and is recruited to 80S ribosomes upon nutrient limiation^12,13^. Rather than causing dimerization, Stm1 and its metazoan orthologue SERBP1 occlude the mRNA-binding channel in both A- and P-site sites thus forming stable inactive 80S particles^12,14^. As expected for a ribosome inactivation factor, when Stm1 is added to yeast translational extracts, it strongly inhibits their activities ^11^.

We reasoned that disrupting the genes encoding for ribosome inactivating factors would yield more reproducible and active bacterial (*Bacillus subtilis Δhpf*) and yeast (*Saccharomyces cerevisiae stm1 Δ*) cell-free translational systems. This could be an especially promising strategy in the case of *B. subtilis.* Since this bacterium already expresses its HPF during exponential growth (although at significantly lower levels than in stationary phase)^8^, it is expected that cell-free translational systems prepared from the Δ*hpf* strain would be more active than those prepared from the wild type strain regardless of growth phase. A popular strategy for preparation of stalled ribosomal complexes utilizes cell-free translation of dicistronic *2Xerm*-mRNA encoding two identical Erm-stalling leader peptides^15^. Stalled ribosomal dimers formed in the presence of the antibiotic erythromycin are readily separated from 70S monosomes – but not from 100S particles – on sucrose gradients. Use of the Δ*hpf* strain lacking 100S ribosomes to generate extracts avoids this additional procedure increasing yields and reducing the hands-on time.

## Results

### Elimination of HPF improves the efficiency of *B. subtilis* coupled transcription-translation system

We opted for a coupled transcription-translation system utilizing the pIVEX2.3MCs FFluc plasmid^16^ that encodes the firefly luciferase ORF preceded by *B. subtilis* optimized ribosome binding site (RBS). Transcription of the mRNA is driven by recombinant T7 RNA polymerase added to the lysate^17^ and the efficiency of protein synthesis was quantified by measuring the luminescence of the translated luciferase protein. For preparing cell-free extracts we used the wild type 168 *B. subtilis* strain and an isogenic Δ*hpf* mutant that displays no growth defect, except for a moderate increase in the lag phase^8^. Our protocol for preparation of a *B. subtilis* transcription-translation system was based on that of Krinsky and colleagues^18^. The system is a binary mixture of cell-free extract and compound mix (**Figure 1A**). The cell-free extract contains a full set of cellular components carrying out protein synthesis, i.e. ribosomes, tRNAs, aminoacyl tRNA synthetases, methionyl-tRNA formyltransferase and translational factors. The compound mix contains i) inorganic ions, importantly Mg^2+^, the key player in ribosomal function^19^ ii) buffering (HEPES) and reducing (DTT) agents iii) NTPs that serve both as the energy source and as the building blocks for mRNA synthesis iv) template DNA in a form of linearized pIVEX2.3MCs FFluc plasmid supplemented with recombinant T7 RNAP polymerase v) amino acids and folinic acid that serve as building blocks for protein synthesis and, finally, vi) stabilizing agents such as PEG-8,000 and glucose.

**Figure 1.**
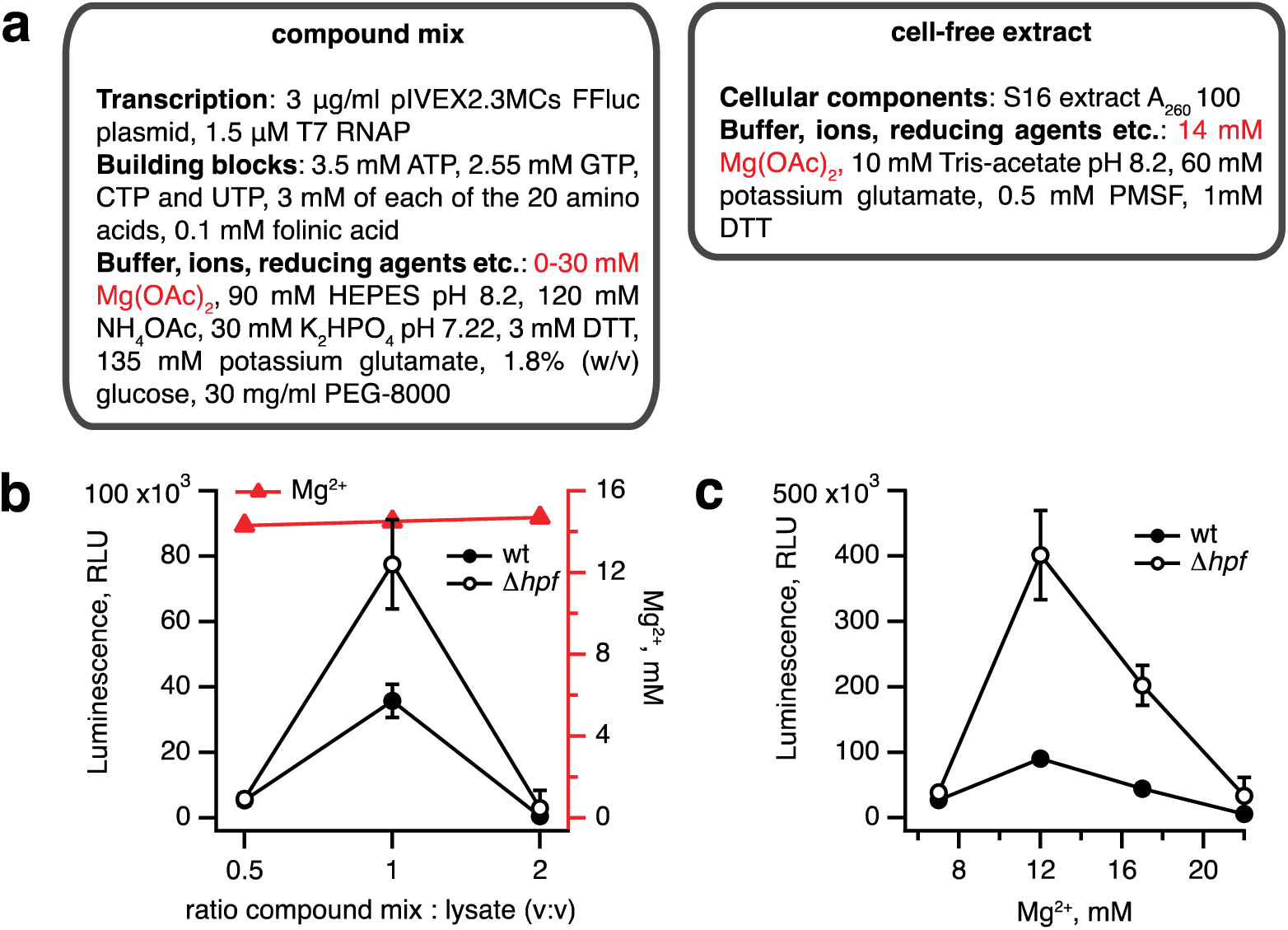
Elimination of HPF improves the efficiency of *B. subtilis* coupled transcription-translation system. (**a**) A cell-free translational system was assembled by combining the compound mix with the cell-free extract. The efficiency of translation was quantified by the activity of the firefly luciferase using the Steady-Glo Luciferase Assay (Promega). Titrations of the compound mix to cell-free extract ratio (**b**) and magnesium ion concentration (**C**) in the cell-free translational system. Luminescence readings were taken after incubation for 1 hour at 37°C. Error bars indicate the standard error of the geometric mean of biological replicates (n ≥ 3).

To achieve a high efficiency transcription-translation system requires sequential optimization by titrating key components. As a first step, we have varied the ratio between the compound mix and cell-free extract (**Figure 1B**). At a 1:1 ratio the activity is optimal, and the Δ*hpf* lysate is approximately two-fold more active. The second optimization step was finding the optimal concentration of magnesium ions^20^. Maintaining the ratio between the extract and compound mix at 1:1, we titrated the final concentration of Mg^2+^ from 7 to 22 mM (**Figure 1C**). While the activity of the Δ*hpf* lysate peaks at 12 mM Mg^2+^, reaching an excess of 400,000 relative light units (RLU), the Δ*hpf* lysate is more active than the wild type at all magnesium concentrations tested.

### Elimination of Stm1 improves the efficiency of *S. cerevisiae* translation system

The *S. cerevisiae stm1*Δ strain was constructed by deleting the *STM1* gene in the wild type MBS^21^ strain. We have opted for a translational system supplemented with an *in vitro* transcribed, capped polyadenylated luciferase mRNA. Capping mRNA dramatically increases the efficiency of translation ^22^ but cannot be performed *in situ* in the lysate and must be added enzymatically to the mRNA after transcription. The translation protocol was based on that of Sachs and colleagues^1^. Just as the bacterial coupled translation-transcription system, the yeast translation system is also a binary mixture of cell-free extract and a compound mix (**Figure 2A**). Since transcription is performed separately, the compound mix contains only the two NTP species necessary for translation, i.e. GTP and ATP, and mRNA is stabilized by RNase inhibitor (rRNasin). Creatine phosphokinase and phosphocreatine serve an energy recuperation system.

**Figure 2.**
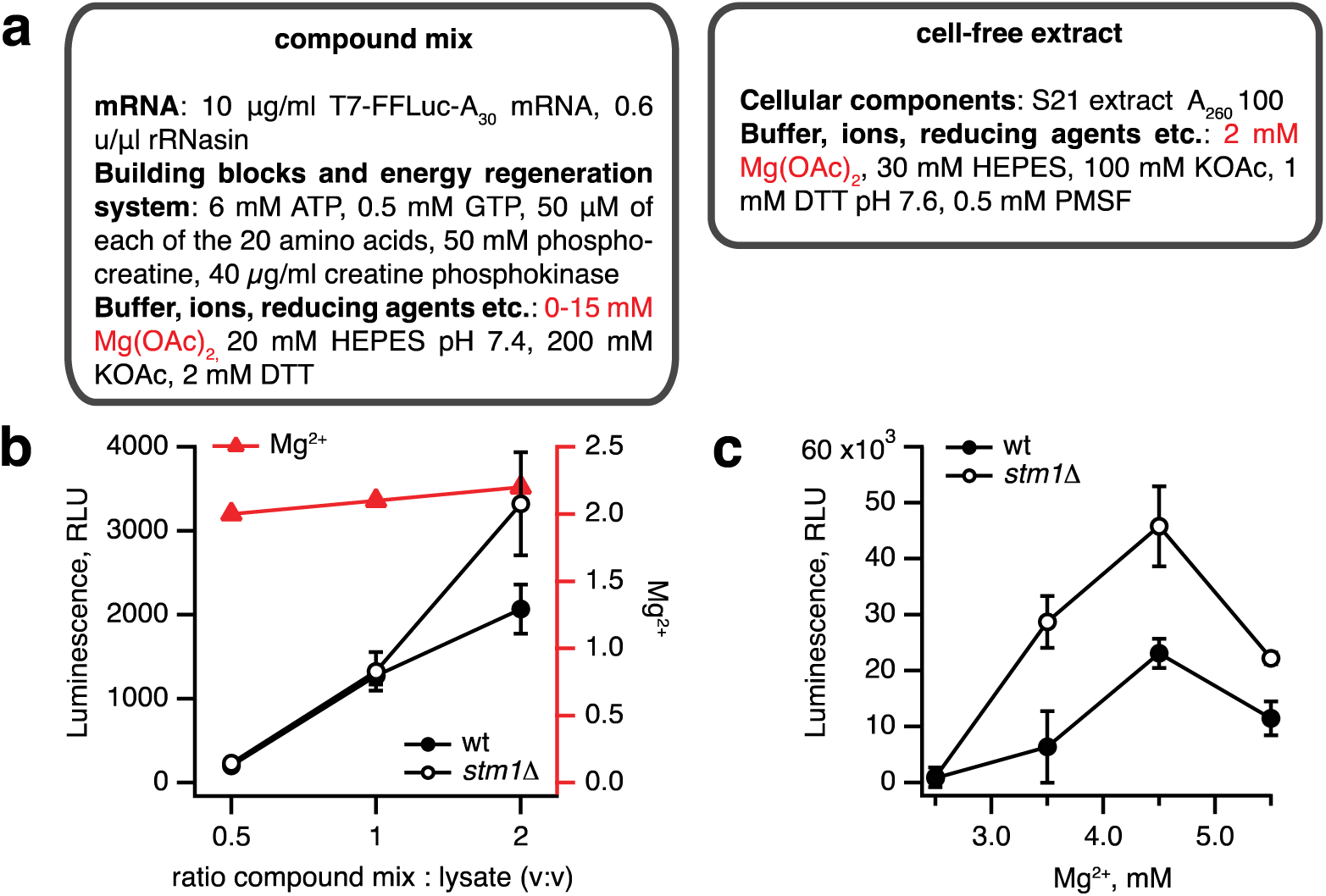
Elimination of Stm1 improves the efficiency of *S. cerevisiae* translation system. (**a**) A cell-free translational system was assembled by combining the compound mix with the cell-free extract. The efficiency of translation was quantified by the activity of firefly luciferase using the Steady-Glo Luciferase Assay (Promega). Titrations of the compound mix to cell-free extract ratio (**b**) and magnesium ion concentration (**c**) in the cell-free translational system. Luminescence readings were taken after incubation for 1 hour at 25°C. Error bars indicate the standard error of the geometric mean of biological replicates (n ≥ 3).

Similarly to the *B. subtilis* system, we have performed two titrations: altering the ratio between the compound mix and cellular extract (**Figure 2B**) and the amount of Mg^2+^ in the extract (**Figure 2C**). The efficiency of the system gradually increases with the percentage of the compound mix in the reaction from 33 to 66% (**Figure 2B**). Notably, both at 33 to 50% compound mix there is no significant difference between the wild type and *stm1*Δ lysates; only in the presence of 66% compound mix *stm1*Δ does the translation reaction display higher activity. However, even in this case, the activity is two orders of magnitude lower than that seen in the *B. subtilis* system. A likely culprit is a sub-optimal concentration of Mg^2+^ ions; as the fraction of the compound mix increases, the Mg^2+^ concentration is moderately increased from 2 to 2.2 mM, following the concomitant increase in activity (**Figure 2B**). Therefore, we titrated Mg^2+^ from 2.5. to 5.5 mM (**Figure 2C**). The activity peaked at 4.5 mM Mg^2^+, reaching an excess of 40,000 RLU with the *stm1*Δ *S. cerevisiae* lysate robustly outperforming the wild type lysate.

## Discussion

Here we demonstrated that genomic disruption of genes encoding ribosome inactivating factors – HPF in *B. subtilis* and Stm1 in *S. cerevisiae* – moderately but robustly improves the activity of bacterial and yeast translational systems. The effect is modest compared with effects of reducing cellular proteolytic or RNase activities: a well-proven approach for genetic manipulation of strains for *in vitro* translation, as exemplified by the *B. subtilis* WB800N strain lacking eight protease-encoding genes^23^ that is used for cell-free translation^24^ and the *E. coli* MRE600 strain with low RNase I activity^25^ that is used for preparation of active ribosomes^26^ and translational systems^27^. In the case of *B. subtilis* WB800N, the cell-free translational system prepared from this strain is 72 times more active than that made from the wild type^24^ – a dramatic result compared to a two-fold effect observed upon *hpf* deletion. A possible next step in developing strains for more efficient cell-free translation system is by combining genetic modifications targeting proteolysis, RNA degradation and ribosome hibernation.

## Methods

### Preparation of 6His-tagged recombinant T7 RNA polymerase (T7 RNAP)

*E. coli* BL21 strain (NEB) lacking the λDE3 lysogen was transformed with pQE30-T7RNAP plasmid (Amp)^17^. An overnight culture in LB media supplemented with 50 μg/mL of ampicillin was used to inoculate a large-scale culture in the same media grown at 37°C with shaking. At OD_600_ of 0.5-0.6 expression of 6His-tagged T7 RNAP was induced by addition of IPTG to final concentration of 1 mM. After an additional two hours growth, cells were harvested by centrifugation, resuspended in buffer A (150 mM NaCl, 100 mM Tris:HCl pH 7.5, 2 mM MgCl_2_, 1 mM β-mercaptoethanol) supplemented with 0.1 mM PMSF, 35 μ.g/mL lysozyme and 1 u/mL DNase I and lysed by one pass via Stansted Fluid Power SFPH-10 Stansted Pressure Cell/Homogenizer (1.5 bar). After removal of the cell debris by centrifugation (35,000 rpm for 40’), the clarified lysate was loaded onto 1 mL HisTRAP HP column (GE Healthcare) equilibrated in buffer A. The column was washed with high salt buffer B (same as buffer A except for 2 M NaCl), and the protein was eluted by a gradient of buffer C (buffer A supplemented with 0.5 M imidazole), pure fractions were pooled, concentrated and buffer-exchanged into storage buffer (100 mM KCl, 20 mM Tris:HCl pH 7.5, 5 mM MgCl_2_, 1 mM DTT, 50% glycerol) using 50 MWCO centricons (Amicon). The purity of protein preparations was assessed by SDS PAGE and spectrophotometrically (A_280/260_ ratio of approximately 1.8). The protein was stored at -20°C.

### Preparation of firefly luciferase mRNA for use in yeast translational lysates

Firefly luciferase mRNA containing a 30 nucleotide poly(A) tail was *in vitro* transcribed from the luciferase T7 control plasmid (Promega) linearized by Afe I as a DNA template for the T7 RNA Polymerase (HiScribe^™^ T7 High Yield) RNA Synthesis Kit. A typical 20 μl reaction containing 1 pmole DNA was incubated for 2 hours at 37°C prior to mRNA isolation with MEGAclear^™^ Kit (Ambion), followed by capping by the Vaccinia Capping System (NEB) and re-purification of mRNA with MEGAclear^™^ Kit (Ambion). The quality of the final product was confirmed by denaturing agarose electrophoresis.

### *B. subtilis and S. cerevisiae* strains

The wild type 168 (*trpC2*) *B. subtilis* strain was provided by Yuzuru Tozawa and the isogenic Δ*hpf* knockout *B. subtilis* strain RIK2508 (*trpC2* Δ*hpf*)) was provided by Fujio Kawamura^8^.

A *S. cerevisiae* strain deleted for *STM1* (MJY1079, *MATa ura3*-*1 leu2*-*3,112 his3*-*11*,*15 trp1*-*1 ade2*-*1 can1*-*100 stm1::HIS3MX6* L-o M-o) was constructed by transforming the MBS strain^21^ with a *stm1::HIS3MX6* DNA fragment PCR amplified from pFA6a-*HIS3MX6*^28^. The oligonucleotides used were: 5’-AGTAGAAATAAACCAAGAAAGCATACACATTTTATTCTCACGGATCCCCGGGTTAATTAA-3’ and 5’-TTATTGGATTCTTTCAGTTGGAATTATTCATATATAAGGCGAATTCGAGCTCGTTTAAAC-3’. The deletion was confirmed by PCR using primers that annealed outside of sequences present in the transformed DNA fragment.

### Preparation of *B. subtilis* bacterial cell-free extract

The lysate preparation protocol is based on that of Krinsky and colleagues^18^ with minor modifications. 50 mL LB cultures of *B. subtilis* 168 wild type and Δ*hpf* strains were inoculated with single colonies from fresh LB plates and grown at 37°C with vigorous shaking overnight. To generate the biomass, two 2L flasks containing 800 mL LB were inoculated to a starting OD_600_ of approximately 0.05 and bacteria were grown at 37°C with shaking. At the OD_600_ of 1.8-2.2 cells were collected by centrifugation at 10,000*g* for 3 minutes (4°C, Beckman JLA-10.500 rotor), pellets dissolved in 100 mL of ice-cold lysis buffer (10 mM Tris-acetate pH 8.0, 60 mM potassium glutamate, 14 mM magnesium acetate, 0.5 mM PMSF, 1mM DTT, pH 8.2), pelleted (3’ at 10,000*g*, 4°C), taken up in 50 mL lysis buffer, and pelleted again in 50 mL Eppendorf centrifuge tubes (30’ at 3,000*g* 4°C) yielding 4-5 grams of biomass that was processed directly. To lyse the cells, lysis buffer was added to 4-5 grams of cells to final volume of 12 mL, and cells were passed once though Stansted Fluid Power SFPH-10 Stansted Pressure Cell/Homogenizer at 2 bar. The lysate was clarified (10’ at 16,000*g*, 4°C) and the supernatant was then desalted using Zeba Spin Desalting Columns 5 mL (ThermoFisher) equilibrated with lysis buffer. After adjusting A_260_ to 100 absorbance units with the lysis buffer, lysates were aliquoted in 50-100 μL fractions, snap frozen in liquid nitrogen and stored at -80°C.

### Preparation of *S. cerevisiae* cell-free extract

The lysate preparation protocol is based on that of Sachs and colleagues^1^ with minor modifications. The wild type MBS and MJY1079 strains were grown in YPD medium at 30°C for 24 hours. To generate the biomass two 2L flasks containing 800 mL YPD were inoculated to a starting OD_600_ of approximately 0.001 and yeast were grown overnight at 30°C until OD_600_ of 4.0-7.0. Cells were collected by centrifugation (10’ at 7,000*g*, 4°C), washed (3’ at 10,000*g*, 4°C) three times with 50 mL of ice-cold lysis buffer (30 mM HEPES, 100 mM potassium acetate, 2 mM magnesium acetate, 0.5 mM PMSF, 1 mM DTT, pH 7.6), and pelleted in 50 mL Falcon centrifuge tubes (30’ at 3,000*g*, 4°C) yielding 10 grams of cells. Cells were frozen as small pellets by mixing with lysis buffer and dropped into liquid nitrogen and stored at -80°C or processed directly. To lyse the cells, 10 g of frozen cells were combined with 1 mL of frozen lysis buffer and crushed with a mortar and pestle in liquid nitrogen for 20 minutes. Note that more robust lysis methods have been shown to reduce the ability to translate exogenous mRNA in a yeast *in vitro* translation system^29^. The resulting lysate was transferred to a 50 mL falcon tube and incubated on ice until melted. The lysate was clarified (30’ at 3,000g followed by ultracentrifugation of the supernatant for 10’ at 21,000g, all at 4°C). The supernatant was desalted using Zeba Spin Desalting Columns 5 mL (ThermoFisher) equilibrated with lysis buffer. After adjusting A_260_ to 100 absorbance units with the lysis buffer, lysates were aliquoted in 50-100 μl fractions, snap frozen in liquid nitrogen and stored at -80°C.

### Preparation of *B. subtilis* transcription-translation system: final optimized protocol

The protocol was based on that of Krinsky and colleagues^18^, with minor modifications. The translation system was assembled by mixing the *B. subtilis* lysate (see above) with the compound mix (10 mM Mg(OAc)_2_, 90 mM HEPES pH 8.2, 30 mM K_2_HPO_4_ pH 7.22, 135 mM potassium glutamate, 120 mM NH_4_OAc, 30 mg/mL PEG-8,000, 1.8 % (w/v) glucose, 3 mM DTT, 3 mM each of the 20 amino acid, 0.1 mM folinic acid, 3.5 mM ATP, 2.55 mM of GTP, CTP and UTP, 3 μg/mL plasmid pIVEX2.3MCs FFluc linearized by Afe I and 1.5 μM recombinant T7 RNAP) to a final volume of 30 μl per reaction point. The lysate and compound mix were combined at a 1:1 ratio. After gently mixing the binary system by pipetting, the reaction was incubated at 37°C for 1 hour with shaking (500 rpm), and 10 μL of the reaction were added to 50 μL of Steady-Glo Luciferase Assay (Promega) (see below).

### Preparation of *S. cerevisiae* translation system: final optimized protocol

The protocol was based on that of Sachs and colleagues^1^ with minor modifications. The translation system was assembled by mixing the yeast lysate (see above) with the compound mix (7 mM Mg(OAc)_2_, 20 mM HEPES pH 7.4, 200 mM KOAc, 2 mM DTT pH 7.4, 50 μM each of the 20 amino acids, 6 mM ATP, 0.5 mM GTP, 50 mM phosphocreatine, 40 μg/mL creatine phosphokinase, 10 μg/mL capped firefly mRNA with a 30 nucleotide poly(A) tail and 0.6 u/μL rRNasin) to a final volume of 30 μl per reaction. Prior to use, the lysates were treated with micrococcal nuclease (NEB) (24 units/μl) activated by 0.5 mM CaCl_2_ (final concentration) at room temperature in order to degrade cellular mRNA species. After a 20’ treatment with the nuclease, the reaction was stopped by addition of EGTA (pH 7.5) to a final concentration of 2 mM. Capped luciferase mRNA was refolded (7’ at 70°C and kept on ice prior to use) and added last to the compound mix immediately prior to assembling the final reaction mixture. The lysate and compound mix were combined to a 1:1 ratio. After gently mixing the binary system by pipetting, the reaction was incubated at 25°C for 1 hour on an Eppendorf Thermomixer with shaking (500 rpm), and 10 μl of the reaction were added to 50 μl of Steady-Glo Luciferase Assay (Promega) (see below).

### Luciferase assay

Steady-Glo Luciferase Assay (Promega) was used as per the manufacturer’s manual. The luciferase assay reagent was aliquoted in 50 μl fractions in 1.5 mL Eppendorf tubes kept in dark at room temperature. Readings were taken after addition of 10 μl reaction mixture to pre-aliquoted luciferase reagent using GloMax 20/20 Luminometer (Promega).

### Data availability

No datasets were generated or analyzed during the current study.

## Acknowledgments

We are grateful to Yuzuru Tozawa for sharing wild type 168 (trpC2) *B. subtilis* strain and Fujio Kawamura for providing the isogenic Δ*hpf* knockout (trpC)^8^, Aivar Liiv for providing pQE30-T7RNAP plasmid^17^, Allan Jacobson for providing the MBS strain^21^ and to Daniel N. Wilson for providing pIVEX2.3MCs FFluc plasmid^16^. This work was supported by the funds from European Regional Development Fund through the Centre of Excellence for Molecular Cell Technology (VH); the Molecular Infection Medicine Sweden (MIMS) (VH); Swedish Research council (grant 2013–4680 to VH and grant 2017-04663 to TN); Ragnar Söderberg foundation (VH); Magnus Bergvalls Foundation (2017-02098 to MJ), Åke Wibergs Foundation (M14-0207 to MJ).

## Author contributions statement

VM and VH conceived and coordinated the study. TB, VM and VH drafted the manuscript together with input from MJ, TN and HT. VM, TB and VH designed experiments and analyzed the data. TB, VM, MJ and HT performed experiments. All authors have read and approved the manuscript as submitted.

## Additional information

### Competing interests

The authors declare no competing interests.

